## Supplementary Figures for "Turn-On Protein Switches for Controlling Actin Binding in Cells"

**a**

| | 1 | 10 | 20 |
|---|---|---|---|
| Lifeact | MGVADLIKKF | ESISKEE | |
| ZO-1 ABS | PAMKPQSVLT | RVKMFENKRS | ASLETKKD |

**b**

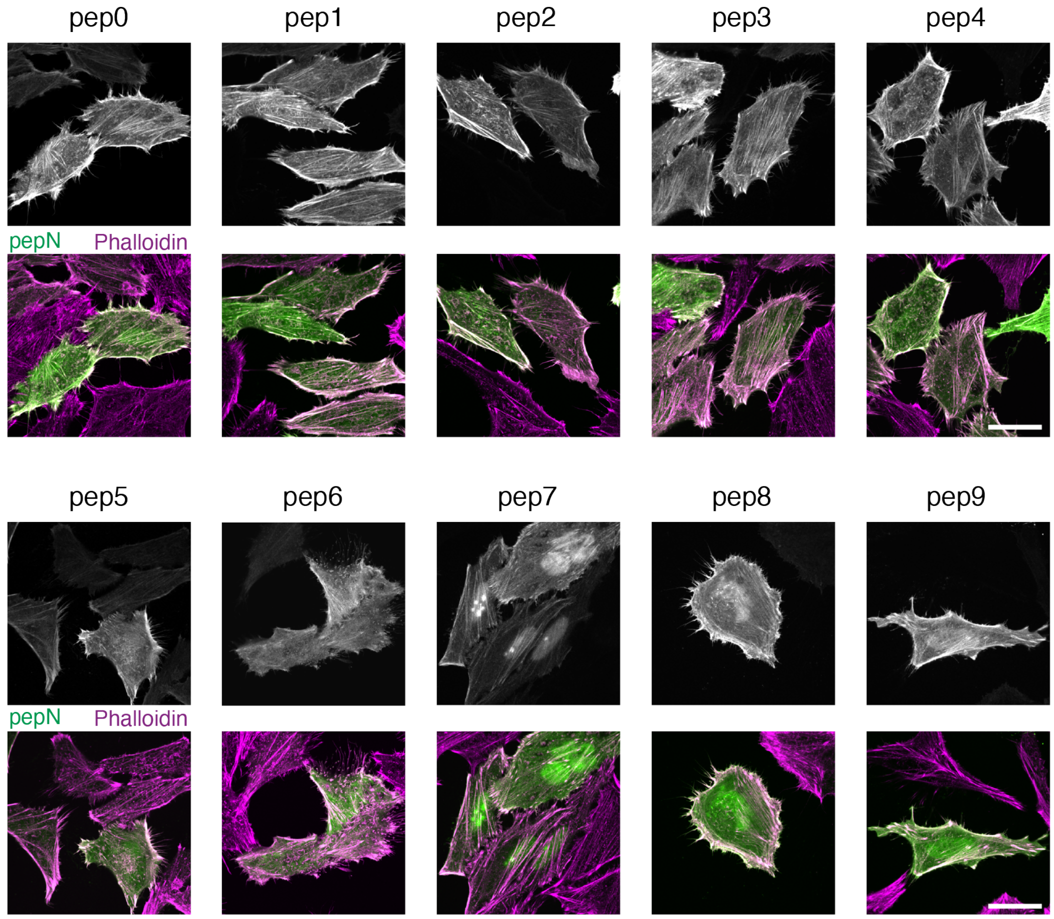


**Supplementary Figure 1. Visualization of ABS-based pepCAST candidates in cells.**
**a,** Amino acid sequences of the ABMs, ZO-1's ABS and Lifeact.
**b,** Fluorescent micrographs of fixed HeLa cells expressing ABS pep0-pep9 in the absence of SZ21. Total cellular actin was visualized by Phalloidin staining (magenta). Scale bar, 30 μm.

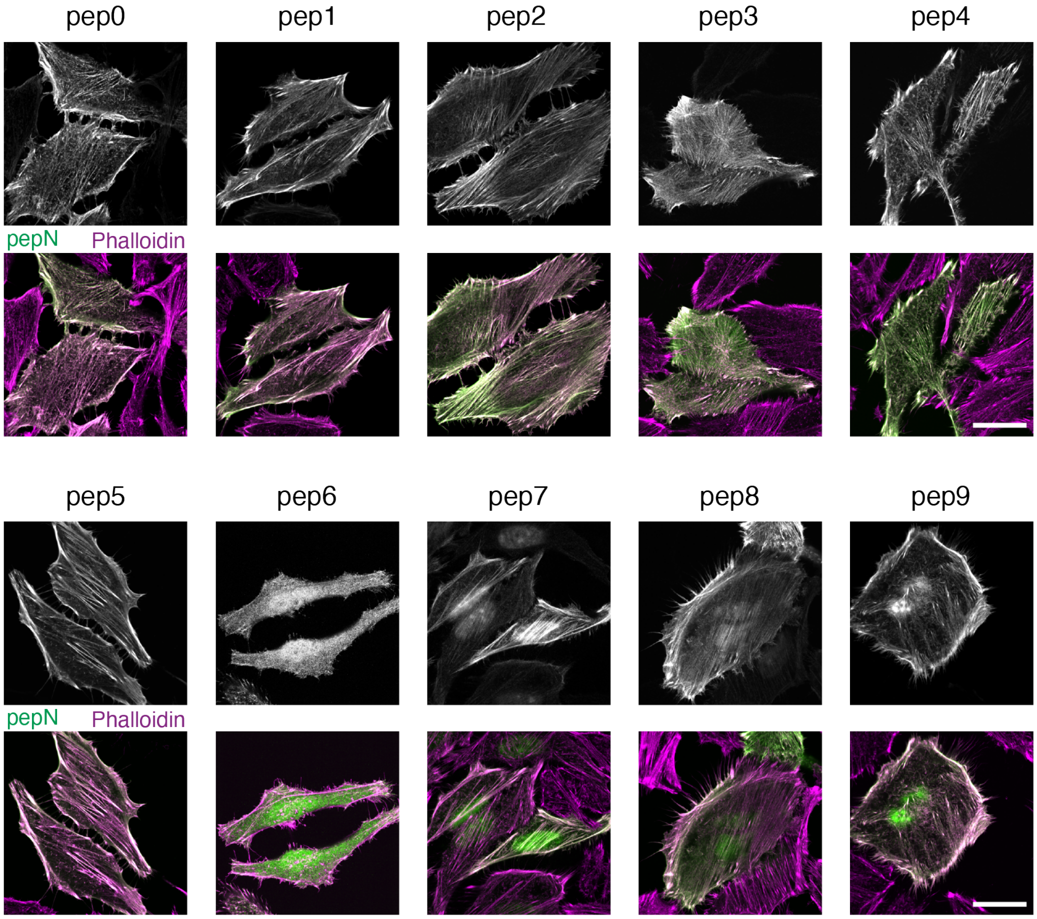


**Supplementary Figure 2. Visualization of Lifeact pep CAST candidates in cells.** Fluorescent micrographs of fixed HeLa cells expressing Lifeact pep0-pep9 in the absence of SZ21. Total cellular actin was visualized by Phalloidin staining (magenta). Scale bar, 30 µm.

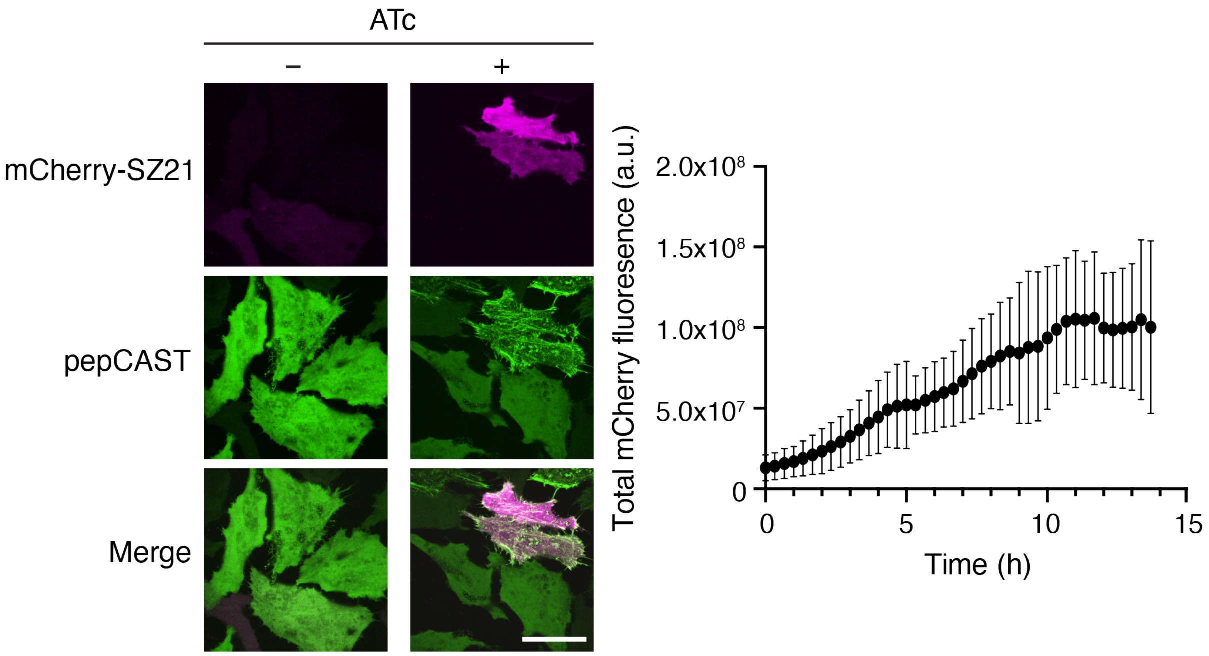


**Supplementary Figure 3. Characterization of SZ21 expression after induction.** Fluorescent micrographs of live stable pepCAST-expressing HeLa cells transfected with mCherry-SZ21 in a TET inducible plasmid in the absence or presence of 1 μM ATc (left). Fluorescence intensity of mCherry shows increasing SZ21 expression in cells over time following the addition of 1 μM ATc (right). Scale bar, 30 μm.

**a**

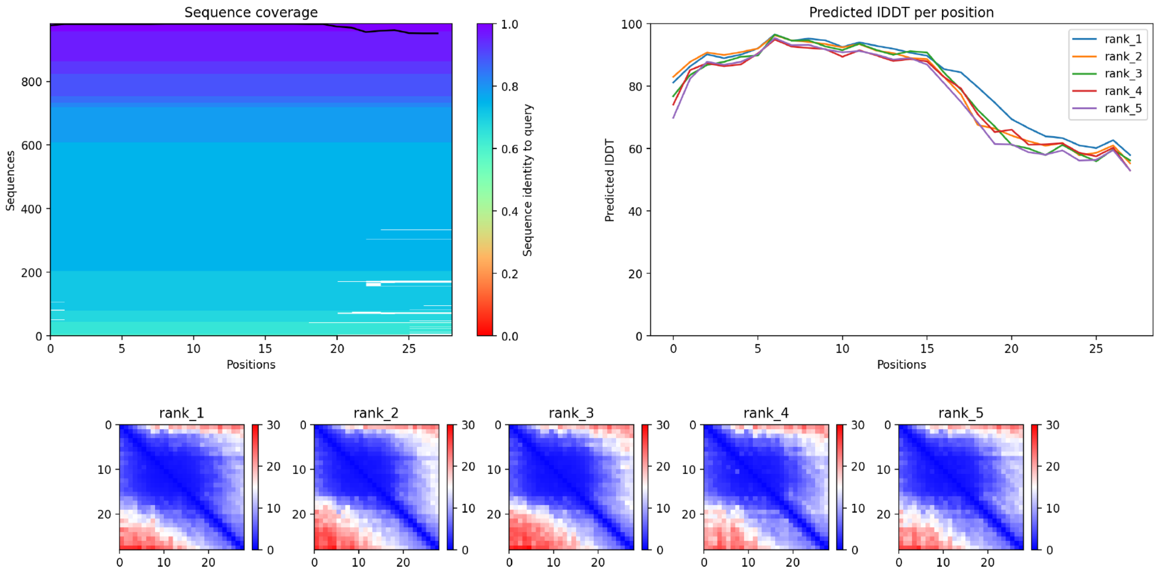


**b**

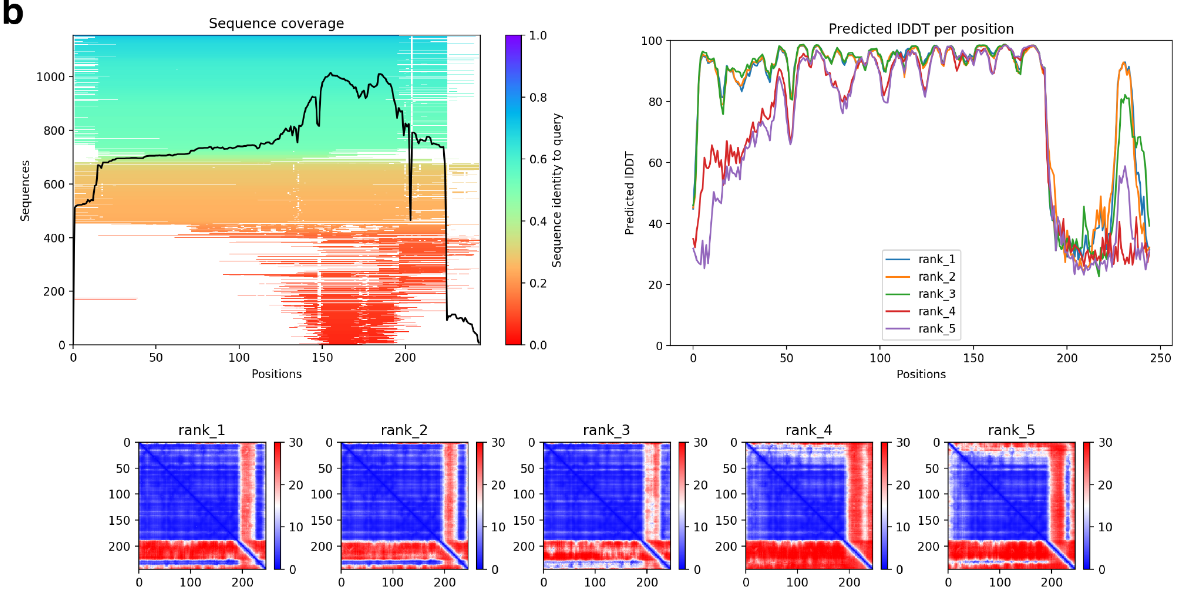


**Supplementary Figure 4. AlphaFold2 structure prediction metrics.**
**a,** Confidence measures for predicted ZO-1 ABS structure showing sequence coverage per residue (top panel, left), local Distance Difference Test (IDDT) scoring (top panel, right), and predicted alignment error (PAE) scoring (bottom panel).
**b,** Confidence measures for predicted smCAST structure showing sequence coverage per residue (top panel, left), local Distance Difference Test (IDDT) scoring (top panel, right), and predicted alignment error (PAE) scoring (bottom panel).

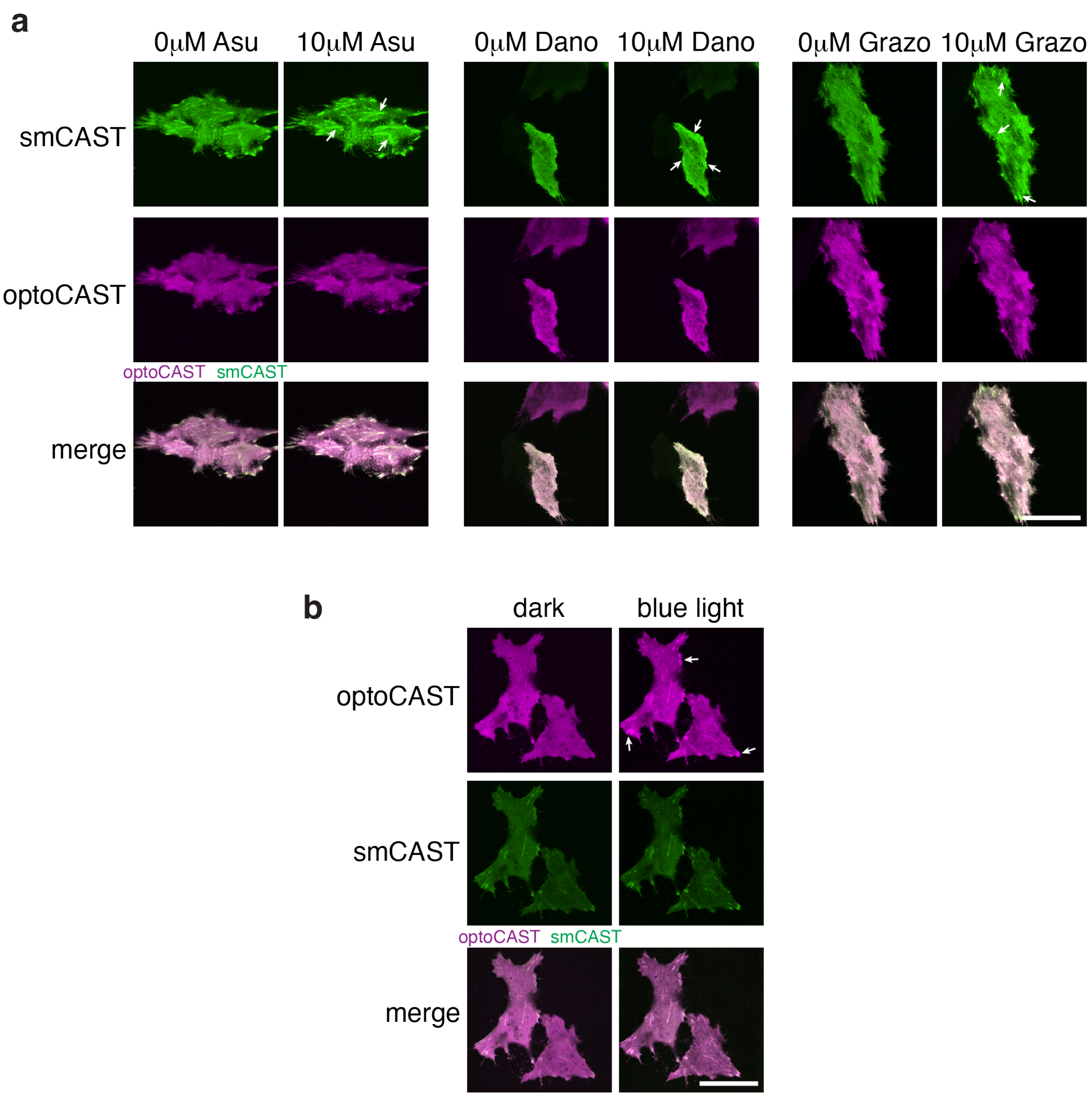


**Supplementary Figure 5. Orthogonal activation of smCAST and optoCAST in cells.**
**a,** Fluorescent micrographs of live HeLa cells co-expressing smCAST and Lifeact optoCAST in the absence or presence of 10 μm Asu, Dano, or Grazo. Only smCAST responds to the small molecule inhibitors and binds to F-actin (white arrows). Scale bar, 30 μm.
**b,** Fluorescent micrographs of live HeLa cells co-expressing smCAST and Lifeact optoCAST in the absence or presence of blue light. Only optoCAST responds to photoactivation and binds to F-actin (white arrows). Scale bar, 30 μm.

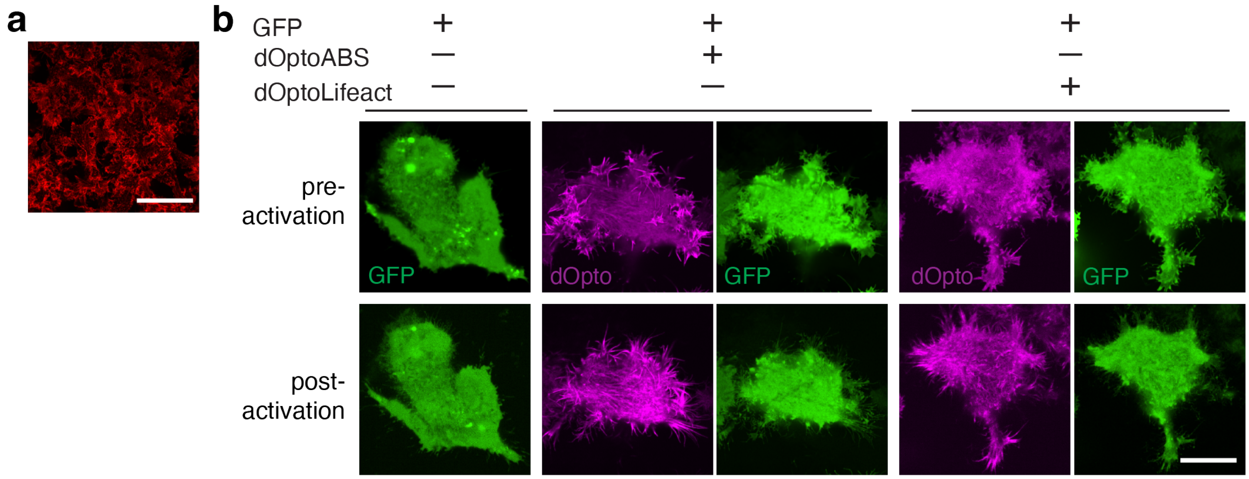


**Supplementary Figure 6. Visualizing cell area in cells expressing dimeric optoCAST.**
**a,** Fluorescent micrograph of HEK 293T cells fixed and stained for endogenous F-actin using Phalloidin. Scale bar, 30 µm.
**b,** Fluorescent micrographs of live HEK 293T cells co-expressing dOptoABS or dOptoLifeact and GFP before and after photoactivation for 10 min. The GFP channel was used for cell area analysis pre- and post-activation. Scale bar, 30 µm.

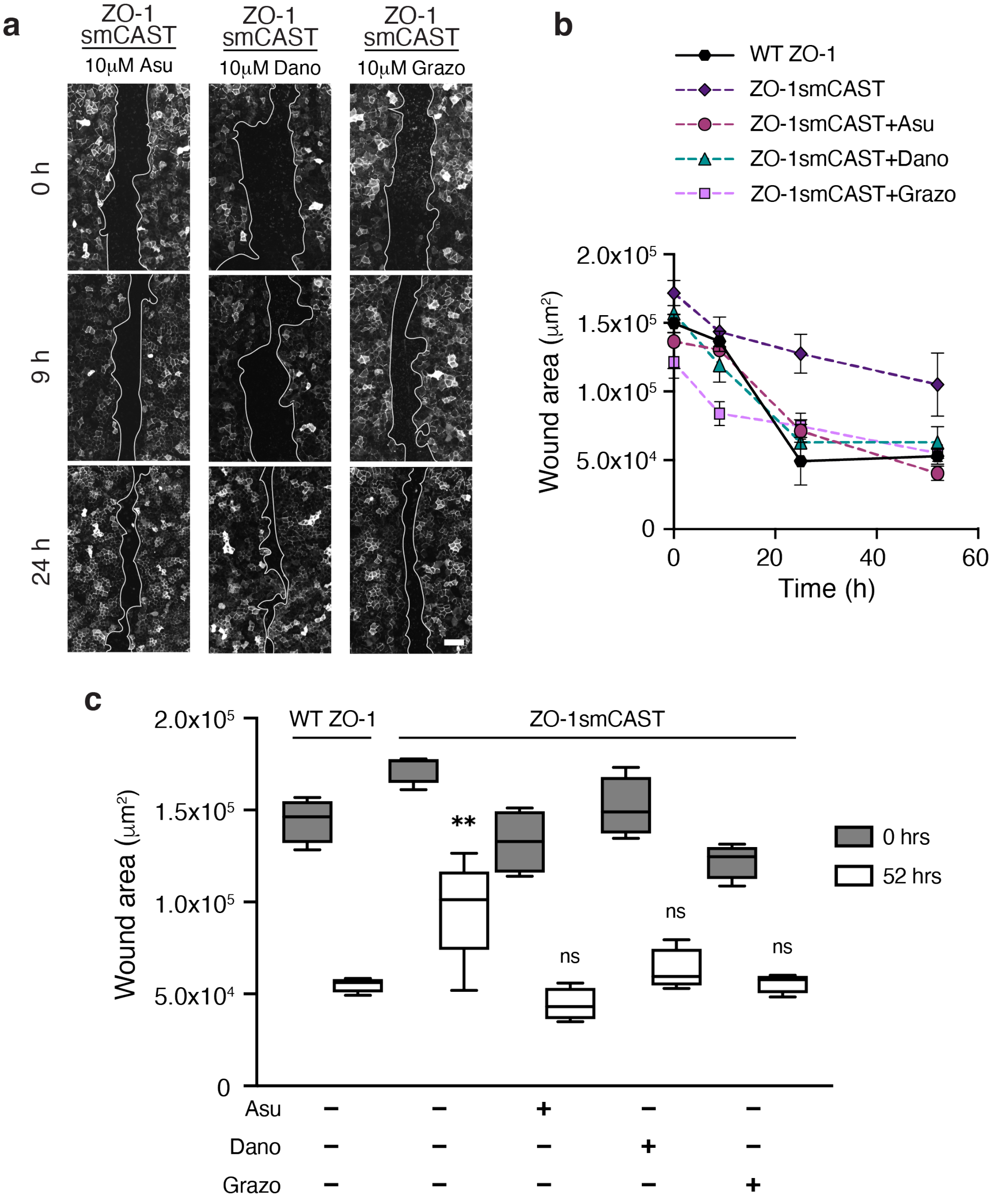


**Supplementary Figure 7. Analysis of collective cell migration with cells expressing CAST-engineered ZO-1.**

**a,** Fluorescent micrographs of collective cell migration in a wounded MDCK II monolayer over time. Stable cell lines lacking ZO proteins and expressing either WT ZO-1 or ZO-1smCAST were imaged following the addition of 10 μm Asu, Dano, or Grazo. Scale bar, 100 μm.

**b,** Quantification of wound area over time. Wound area decreases for cells expressing either WT ZO-1 or ZO-1smCAST and cultured in the absence or presence of 10 μm Asu, Dano, or Grazo.

**c,** Quantification of wound area after 52 h. Box represents 25th to 75th percentiles with the middle line as the median and whiskers as the maximum and minimum values. n = 4 replicates. P-values were determined using an unpaired t-test. (ns, not significant $P > 0.05$; $^{*}P < 0.05$; $^{**}P < 0.01$; $^{****}P < 0.0001$).
