## Supplementary Videos for "Turn-On Protein Switches for Controlling Actin Binding in Cells"

**Supplementary Video 1**
**pepCAST activation following induction of SZ21 expression**. Confocal imaging of a live HeLa cell stably expressing ABS pepCAST and transfected with inducible SZ21. Images were acquired every 20 minutes for 5.5 hours after a 2-hour incubation period of 1 µM ATc to induce SZ21 expression.

**Supplementary Video 2**
**smCAST activation following addition of small molecule, Grazo**. Confocal imaging of a live HeLa cell expressing smCAST. Images were acquired every 5 minutes for 1.5 hours immediately after the addition of 10 µM Grazo.

**Supplementary Video 3**
**optoCAST photoactivation with blue light**. Confocal imaging of a live HeLa cell expressing Lifeact optoCAST. Blue light was pulsed every 2.5 seconds for 6 minutes and images were acquired following each pulse.

**Supplementary Video 4**
**Photoactivation of dimeric optoCAST in single cell**. Confocal imaging of live HEK 293T cells expressing dOptoLifeact. Blue light was pulsed every 2.5 seconds for 8 minutes and images were acquired following each pulse.

**Supplementary Video 5**
**Photoactivation of dimeric optoCAST in tissue island**. Confocal imaging of live MDCK II cells expressing dOptoABS. Blue light was pulsed every 2.5 seconds for 8 minutes and images acquired following each pulse.

**Supplementary Video 6**
**Activation and relocalization of ZO-1smCAST**. Confocal imaging of live HeLa cells expressing ZO-1smCAST. Images were acquired every 5 minutes for 5.5 hours following a 1-hour incubation of 10 µM Dano.
